# Investigating the determinants of performance in machine learning for protein fitness prediction

**DOI:** 10.1101/2020.09.30.319780

**Authors:** Mahakaran Sandhu, Adam C. Mater, Dana S. Matthews, Matthew A. Spence, Artem A. Lenskiy, Colin Jackson

**Affiliations:** Research School of Chemistry, The Australian National University, Canberra, ACT 2601, Australia; ARC Centre of Excellence for Innovations in Peptide & Protein Science, Research School of Chemistry, The Australian National University, Canberra, ACT 2601, Australia; School of Engineering and Technology, The University of New South Wales, ACT 2600, Australia; ARC Centre of Excellence in Synthetic Biology, Research School of Biology, The Australian National University, Canberra, ACT 2601, Australia

## Abstract

Machine learning (ML) has revolutionized protein biology, solving long-standing problems in protein folding, scaffold generation and function design tasks. A range of architectures have shown success on supervised protein fitness prediction tasks. Nevertheless, in the absence of rational approaches for evaluating which architectures are optimal for specific datasets and engineering tasks, architecture choice remains challenging. Here, we propose a framework for investigating the determinants of success for a range of ML architectures. Using simulated (the NK model) and empirical fitness landscapes, we measure sequence-fitness prediction along six key performance metrics: interpolation within the training domain, extrapolation outside the training domain, robustness to increasing epistasis/ruggedness, ability to perform positional extrapolation, robustness to sparse training data, and sensitivity to sequence length. We show that architectural differences between algorithms consistently affect performance against these metrics across both experimental and theoretical landscapes. Moreover, landscape ruggedness emerges as a primary determinant of accuracy of sequence-fitness prediction. Our methodology and results provide a rational strategy for experimental data sampling, model selection and evaluation rooted in fitness landscape theory; one that we hope will advance sequence-fitness prediction accuracy, with implications for protein engineering and variant functional prediction.

## Introduction

In recent years, machine learning has transformed many domains of science, including protein biology and engineering. Models such as AlphaFold have solved decades-long problems in protein structure prediction,^1,2^ and protein language models such as Evolutionary Scale Modelling (ESM) have demonstrated the ability to learn meaningful representations of protein sequences that encode fundamental rules and properties of protein biophysics and evolution.^3–5^ Nevertheless, a key frontier remains: accurate prediction of protein fitness from sequence,^6^ especially in the context of low data availability.^7^ Successfully addressing this key frontier promises to revolutionize the field, leading to better proteins for industry, science and medicine at a fraction of the cost. Several ML methods have been implemented to address this,^8^ including Gaussian process regression,^9–11^ unsupervised statistical analyses and models,^12,13^ deep neural networks and sequence models.^14–18^ However, scientific ambiguity remains regarding how protein fitness landscapes and ML models interact, hindering our ability to identify and address gaps.

Investigating how ML models learn the mapping from protein sequence to fitness requires careful consideration of the fundamental structure on which sequence-fitness relationships exist: the fitness landscape.^19^ The fitness landscape is a well-known idea in biology, first introduced in the seminal work of Sewall Wright, and applied to proteins by John Maynard Smith.^20,21^ A combinatorially complete fitness landscape for a protein is one where the sequence space for a given length protein (i.e. all possible combinations of amino acids for a given length protein) (**Figure 1a, 1b**) are mapped to a corresponding numerical fitness value (i.e. some measurable biophysical property of the protein such as thermostability, fold enrichment, binding affinity or fluorescence) (**Figure 1c)**.^22–24^ A key characteristic of such landscapes is their ruggedness. Intuitively, where the fitnesses of adjacent sequences are similar, there are smooth changes in fitness as the sequence space is traversed along its one-mutant neighbors, resulting in a smooth or correlated landscape with few local maxima (**Figure 1d**).^25^ Conversely, in rugged or uncorrelated landscapes, adjacent sequences can have sharp changes in fitness (akin to sharp crags and deep chasms in geographical landscapes) with many local maxima (**Figure 1e**), making reliable prediction challenging. Highly rugged landscapes are thought to result from epistasis, a fundamental phenomenon in protein evolution wherein the effect of a mutation is dependent on the context into which it is introduced (i.e. the context-dependence of mutations).^26^ Ruggedness has been measured using various different metrics, including number of local maxima, Dirichlet energy and Fourier decomposition.^27,19^

**Figure 1.**
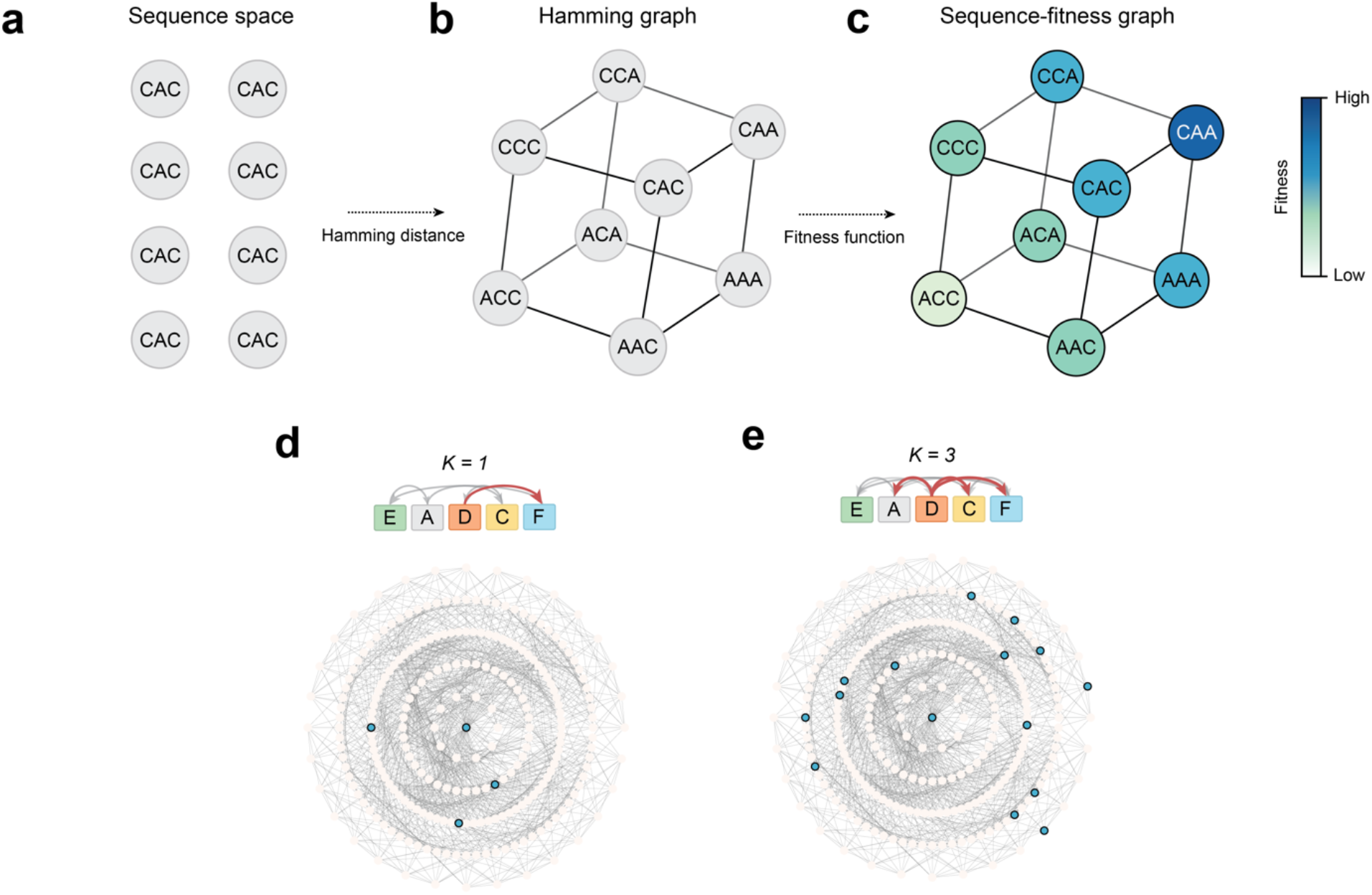
Fitness landscapes and the NK landscape. Depiction of a sequence dataset as a sequence-fitness graph: (**a**) The combinatorial sequence space of an alphabet of two amino acids (A, C) over a sequence length of three. (**b**) This sequence data can then be represented as a Hamming graph, wherein nodes represent sequences and edges connect sequences (nodes) that differ by a single mutation (**c**) each node in the graph maps to a scalar fitness, creating a sequence-fitness graph (which can be thought of as a sequence-fitness *landscape*). (**d**) Schematic of *NK* interactions where *K = 1*: here, the fitness contribution of the *i*^*th*^ position depends on its own identity as well as the identity of, on average, 1 other position. This results in a smoother sequence-fitness graph with few local maxima (four local maxima), here depicted as a radial plot. Each concentric circle of the radial plots is stratified by a mutational regime, with the reference ‘seed’ sequence in the center. This results in a smooth sequence-fitness graph with few local maxima (four local maxima) (**e**) Schematic of NK interactions where *K* = 3: here, the fitness contribution of the *i*^*th*^ position depends on its own identity as well as the identity of, on average, 3 other positions. This results in a rugged sequence-fitness graph with many local maxima (fifteen local maxima).

The goal of ML sequence-fitness prediction is to approximate the fitness function (i.e. the mapping from sequence to fitness), ideally by training on sparse experimental sampling of the fitness landscape. Numerous strategies for experimental sampling exist (e.g. ‘shotgun’ random sampling^28,29^ and sampling along evolutionary trajectories^30–32^). In this work, we use a sampling strategy rooted in fitness landscape theory that allows us to define notions of interpolation within a training dataset regime, and extrapolation away from it (defined below). An important concept to frame the introduction to these sampling strategies is mutational regimes. We consider a mutational regime to be all the sequences in the dataset that differ by *m* number of mutations from an arbitrarily chosen reference sequence (e.g. the wild-type).

Therefore, all the sequences that differ by one mutation will be in the first mutational regime, all those that differ by two mutations in the second mutational regime, and so on.

Here, we sought to understand how fitness landscape ruggedness and training data sampling affect ML performance. To this end, we define a set of key metrics to assess ML sequence-fitness prediction performance: (1) ability to interpolate within the mutational regimes present in the training set; (2) ability to extrapolate beyond the mutational regimes present in the training set; (3) robustness to increasing fitness landscape ruggedness; (4) ability to perform positional extrapolation; (5) robustness to sparse experimental sampling of the fitness landscape; and (6) robustness to increasing sequence length. Formally assessing these metrics requires a single training dataset that exhaustively spans multiple mutational regimes and is of increasing and known ruggedness. The former is prohibitive experimentally, and the latter is subject to numerous ambiguities regarding the measurement and interpretation of fitness landscape ruggedness.

The NK landscape addresses these shortcomings, offering a precise and tunable simulated fitness landscape model rooted in epistasis,^33^ and has previously been used to evaluate ML models^7,34,35^ and in evolutionary analysis.^36,37^ In NK landscapes, the *K* parameter controls the degree of epistasis and hence ruggedness of the landscape, with higher *K* values producing higher degrees of epistasis and ruggedness **(Figure 1d, 1e)**. Indeed, the properties of NK landscapes have been studied in-depth and closed-form expressions have been derived to describe the behavior of evolutionary processes that occur over them.^38^

By simulating landscapes of increasing ruggedness and stratifying sequences into mutational regimes, we assessed a range of sequence-fitness ML models against the six metrics discussed above. We analyzed a diverse variety of traditional and state-of-the-art models (including linear regressors, decision tree models, and a range of neural networks), to show that ruggedness is a key determinant of performance, and that different model architectures differ meaningfully in their abilities to interpolate, extrapolate and learn from sparse data.

## Results

### Generation of synthetic datasets

We first generated simulated protein datasets using the NK landscape model. NK landscapes were generated over a reduced alphabet of 6 amino acids (*A, C, D, E, F, G*) with a sequence length of 6. Using a reduced alphabet and short sequence length permits the resulting combinatorial sequence space to be tractable and amenable to computational modelling and analysis (**SI Table 1**). As the NK landscape is not rooted in a biophysical model, the choice of alphabet size (and indeed the alphabet set itself) is arbitrary. In the NK model, the *K* parameter controls the number of interactions between amino acid sites, with *K* = 0 corresponding to purely additive interactions between adjacent residues (resulting in a smooth fitness landscape) and *K* = 5 corresponding to, on average, 5 interactions per site (i.e. each site interacting with all other sites) (**Figure 1d-e**), yielding a maximally rugged fitness landscape. All modelling and analysis on *NK* landscapes was performed over four replicate landscapes.

### Interpolation and extrapolation performance

On any dataset that spans multiple mutational regimes, interpolation and extrapolation can be assessed concurrently by stratifying data into mutation regimes *M*_*n*_ from some arbitrary seed sequence *M*_0_ (which could be the wild-type), followed by expanding the training data to include an increasing number of mutational regimes (**Figure 2a**). We performed interpolation and extrapolation testing for all models on four replicate NK landscapes of increasing ruggedness (*K* = 0 through to 5 for *N* = 6), starting from randomly chosen seed sequences in each replicate. Their performance on each test set was evaluated as the mean standard error (MSE), Pearson’s correlation coefficient (*r*) and the coefficient of determination (*R*^2^) between the simulated ground-truth values and the model-predicted values.

**Figure 2.**
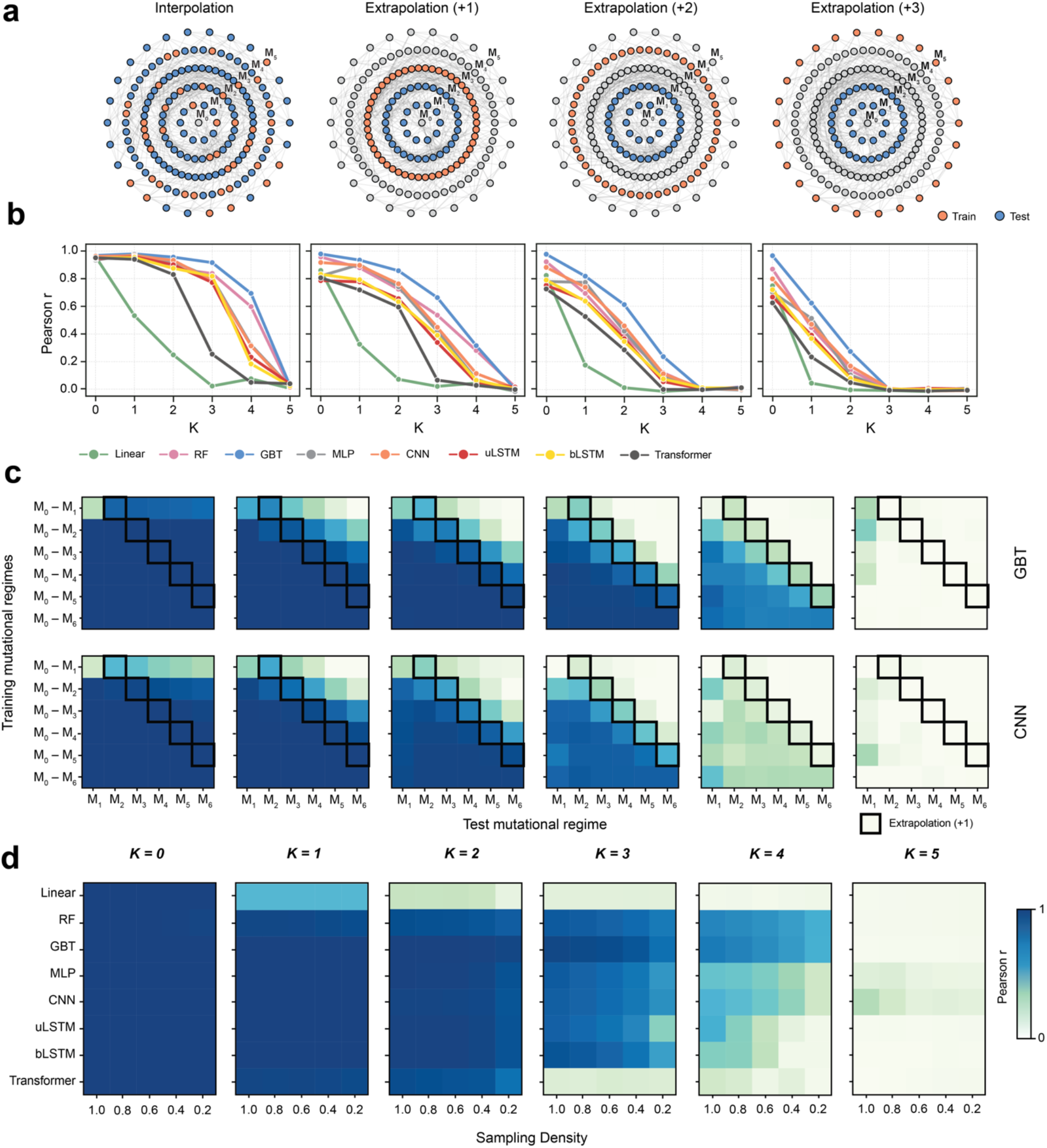
Interpolation, extrapolation and robustness to sparse training data on *NK* landscapes. (**a**) Graphical representation of a complete combinatorial sequence space stratified into mutational regimes *M*_0_ − *M*_5_ from an arbitrarily selected seed sequence (*M*_0_). Specifically, a mutational regime (*M*_*n*_) represents sequences that are *n* mutations away from the seed sequence. In interpolation, both the train and test sets contain sequences from the same mutational regimes, whereas in extrapolation, the test set contains sequences from mutational regimes greater that those in the train set (i.e. mutational regimes *M*_*n*+*m*_ where *m* > 0 and *n* is the highest mutational regime in the train set). In extrapolation, the model is trained on an expanding number of mutational regimes, but performance is always evaluated on mutational regimes greater than those in the train set. (**b**) The performance (Pearson *r*) of models over increasing ruggedness (*K*) on interpolation, extrapolation +1, extrapolation +2 and extrapolation +3 (variants containing one, two or three additional mutations compared to the training data, respectively) test datasets. (**c**) The average Pearson correlation (over 4 replicate NK landscapes) for the GBT and CNN models. Each heatmap shows how the correlation between ground truth and predicted values changes as a larger number of mutational regimes are used for training. Extrapolation into one mutational regime beyond the training data is highlighted in cells with a black border (i.e. variants containing one additional mutation compared to the training data, *M*_*n*+*m*_where *m* = 1). To the right of the cells in the black border are higher extrapolation regimes (i.e. variants containing two to five additional mutations compared to the training data, *M*_*n*+*m*_ where *m >* 1), whereas to the left are interpolation regimes (*M*_*n*+*m*_ where *m* < 1). The y-axis of the heatmaps corresponds to mutational regimes in the train set, and the x-axis to mutational regimes in the test set. (**d**) The performance (Pearson *r*) of all models on *NK* landscapes of increasing ruggedness (*K*) as train data sampling density decreases, measuring robustness to sparse data. The y-axis of these heatmaps corresponds to the models tested, the y-axis to the test set sampling density. Each consecutive heatmap denotes training on incrementally higher *K* value *NK* landscapes (from *K* = 0 to *K* = 5).

All models perform worse on interpolation as ruggedness increases (**Figure 2b-c, SI Figure 1**). At the limit of completely uncorrelated landscapes (*K* = 5 for *N* = 6) all models fail dramatically at both interpolation and extrapolation (**Figure 2b-c, SI Figure 1**). Ability to extrapolate correlates closely and inversely with ruggedness: as ruggedness increases, extrapolation performance decreases. For example, at *K* = 2, the GBT model can perform reasonable positional extrapolation to 3 mutational regimes (extrapolation +3) beyond the training data, at *K* = 4 only to 1 mutational regime (extrapolation +1), and at *K* = 5, it fails completely at extrapolation (**Figure 2b**). The decrease in performance as a result of ruggedness is seen across all models (**SI Figure 2)**.

### Robustness to sparse training data

Assaying protein fitness in the wet-lab can be cost- and labor-intensive. A key attraction of *in silico* methods for protein sequence-fitness prediction is their ability to give accurate results in a fraction of the time and money. Nevertheless, all ML models ultimately require training data from wet-lab experiments. Models that can excel at learning from few experimental samples therefore have an advantage over those that require larger volumes of training data. We therefore sought to test the robustness to sparse training data, to identify models that excel at sequence-fitness prediction despite having few training samples. To test this, we segregated the NK landscape datasets into train and test categories. We then randomly subsampled the train data at decreasing sampling densities, beginning with 1 (all the training data) to 0.2 (only 20% of the training data), and used the subsampled datasets to train models.

All models were then tested on test data (**Figure 2d**). As seen in interpolation and extrapolation, the performance of all models declines as landscape ruggedness increases. Additionally, higher landscape ruggedness values also cause a quicker drop in performance as less training data is sampled, although the magnitude of this decline varies between models. The performance of individual models does not differentiate until *K* = 2 where the linear model begins to fail. At *K* = 4, decision tree models (RF and GBT) maintain the best performance and are most robust to data ablation. However, at *K* = 5, the MLP and CNN models are the best performing.

Generally, the neural network (NN) models performed poorly at lower sampling densities compared to the decision tree models. The stark exception to this is MLP and CNN, which outperform all other models at *K* = 5. Of the NN models, transformers performed most poorly, followed by the LSTM models. A likely explanation for this is the complexity of the LSTMs and transformers: these models have many parameters, requiring commensurately larger amounts of data for appropriate training (**SI Table 3)**.

This illustrates a limitation not only in supervised protein sequence fitness prediction but in supervised ML more broadly. The complexity and large number of parameters of deep learning models often means that large amounts of data are required to train them appropriately. In the setting of protein sequence-fitness prediction, acquiring such large training datasets experimentally can be prohibitively resource intensive. Benchmarking candidate models for robustness to low training data volume permits judicious choice of ML model, thereby reducing the resource burdens of experimental data collection.

### Positional extrapolation

An additional type of extrapolation that can be considered involves tasking the model with predicting the influence of a mutation at a position that has not been altered in the training set. We distinguish this form of *positional* extrapolation from the previously introduced form of mutational regime (MR) extrapolation (i.e. extrapolating to higher mutational regimes). Indeed, positional extrapolation can occur inadvertently when performing MR extrapolation. This effect is particularly pronounced when assessing interpolation on *M*_*1*_, as any dataset splitting of this regime is likely to remove all training examples at some positions in the sequence, thus demanding the model perform positional extrapolation.^39^ A key limitation of the NK landscape here is its lack of biophysical grounding, meaning that positional extrapolation is not possible on NK landscapes. Future work on identifying/developing model fitness landscapes with a grounding in biophysics (such as the generalized/structurally informed NK^7^ and NKp^40^ models) could be useful as alternatives to the NK landscape when assessing the ability of models to perform positional extrapolation. This aligns with recent experimental work that demonstrated that simpler neural network architectures excel at local extrapolation for designing high-fitness proteins, while more sophisticated convolutional models can venture deeper into sequence space but may lose functional specificity – findings that parallel our observations of architecture-dependent extrapolation capabilities.^41^

To highlight the challenge presented by positional extrapolation, we tested all models on NK landscapes by stratifying sequences. To this end, we fixed the reference/seed amino acid at a given site in the training set, trained the model on sequences with that amino acid at that site remaining constant, and tested the model’s ability to predict the mutational effect 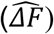 for alternative amino acids at that site **(Figure 3a)**. The predicted mutational effect 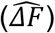 is then compared with the true mutational effect (*ΔF*) to determine the performance of each model at positional extrapolation (**Figure 3a**). We considered this separately from the performance of each model in predicting fitness for the overall sequence, as at low *K* values, a model may be able to roughly predict the fitnesses of variants which contain mutations at sites that were not varied in the training data (positional mutations) by learning the effects of all other sites.

**Figure 3.**
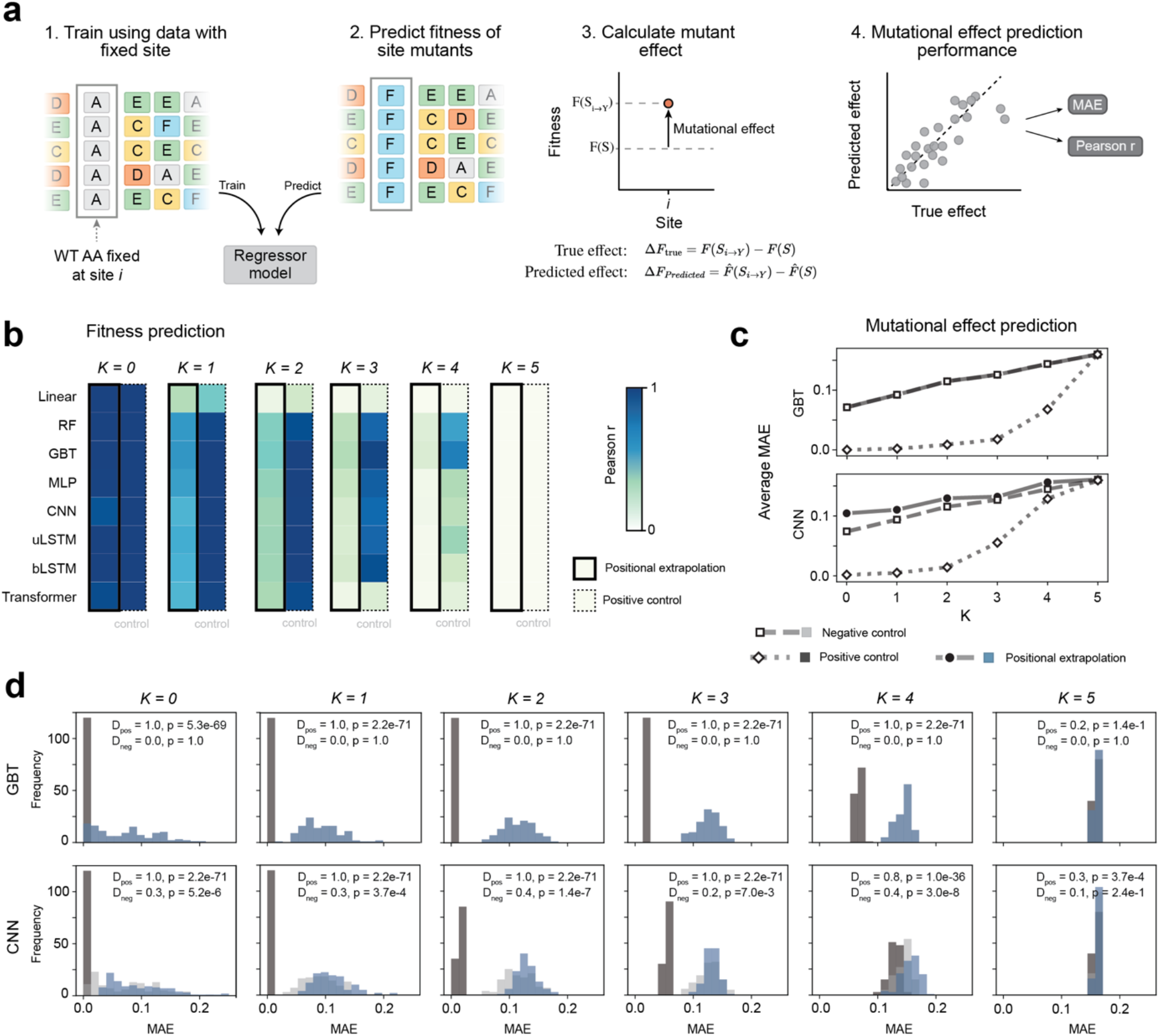
Positional extrapolation on NK Landscapes. (**a**) Schematic for measuring positional extrapolation. Models were trained on sequences with the amino acid at site *i* fixed and tested on sequences with alternative amino acids at site *i* (i.e. amino acids at site *i* that the model was not exposed to in the training set). Mutational effect (*ΔF*) was calculated as the difference in fitness between the sequence with alternative (mutant) amino acids at site *i* (*F*(*S*_*y*→*Y*_)) and the fitness of the sequence with the WT amino acid at site *i* (*F*(*S*)). The predicted mutational effect 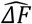 was compared against the true mutational effect *ΔF* to determine the extrapolation performance of each model. (**b**) The Pearson correlation (*r)* between predicted sequence fitness and ground truth sequence fitness. In the positive control, models were exposed to alternative mutant amino acids at the *i*^*th*^ site in training data. (**c**) The average mean absolute error (MAE) of GBT and CNN models in predicting mutational effects (i.e.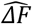) with increasing *K* value. A lower MAE indicates better performance. As in **b**, in the positive control models were exposed to alternative mutant amino acids at the *i*^*th*^ site in training data. In the negative control, NN models were initialized with random weights and were not trained, and decision trees were trained on shuffled targets. The MAE values for the negative control are effectively identical (i.e. follow/trace) to the MAE results for positional extrapolation for the GBT model, but not for the CNN, where positional extrapolation performs better than the negative control. (**d**) Histogram of MAE values for GBT and CNN models when predicting mutational effects 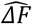 in the positional extrapolation testing regime. *D* denotes Kolmogorov’s *D* statistic. Here, *D*_*neg*,_ denotes the *D* statistic between the distribution of MAE values from positional extrapolation versus the negative control, and *D*_*pos*_ denotes the *D* statistic between the distribution of MAE values from positional extrapolation versus the positive control. Note that D is 1 for sample distributions drawn from separate underlying distributions and 0 for those drawn from the same distribution (i.e. a higher number means there is a statistical difference between distributions).

As the positive control for these experiments, we introduced 80% of the sequences containing alternative amino acids at the *i*^*th*^ site into the training set, with the remaining 20% of these sequences used as the test. Therefore, the positive control establishes model performance when the model has excellent access to information about the effects of alternative amino acids at that site (i.e. when it is not asked to positionally extrapolate). As the negative control, we initialised NN models with random weights without further training and used these untrained models to perform predictions. For the decision tree models, we used shuffled training targets. Hence, the negative control establishes a negative baseline for model performance in which the models have access to no meaningful training information. Comparing positional extrapolation against these negative and positive controls permits us to gauge the relative effectiveness of positional extrapolation.

When considering the performance of each model in predicting the overall fitness of sequences in positional extrapolation, at *K* = 0, there is no difference between this performance and the positive control **(Figure 3b)**. At *K* = 1 however, this performance is reduced compared to the positive control. This trend continues as *K* increases. Interestingly, positional extrapolation performance declines most at *K* = 2 *and K* = 3, where the positive control still maintains good performance. At *K* = 4, all models struggle, with the decision tree models performing poorly but better than the other models. At *K* = 5, both positional extrapolation and the positive control fail dramatically **(Figure 3b)**. This trend is consistent with the fact that, as *K* increases, each position interacts with a larger number of other positions (i.e. higher epistasis); as a result, at high *K* values, a single mutation has an outsized impact on the overall fitness of the sequence. We further evaluated model ability to perform positional extrapolation by considering the mean absolute error (MAE) between the predicted 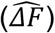 and true mutational effects (*ΔF*). Here, as *K* increases, so too does the MAE of 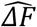 when compared to *ΔF* for both positional extrapolation and the positive control (**Figure 3c, SI Figure 3a**). However, it is apparent that when compared to the negative control, this increase is not necessarily due to positional extrapolation being possible on smoother landscapes, but rather, a difference in the magnitude of effects as *K* increases. This is further highlighted when comparing histograms of the MAEs over increasing *K* values (**Figure 3d**) which shows that performance on positional extrapolation only resembles the positive control (having Kolmogorov D statistics close to 0) at high *K* values, where mutational effects can no longer be predicted.

We note that decision tree models, which are architecturally unable to perform positional extrapolation (being unable to produce nodes for sites with no variance) have similar performances to other models (**Figure 3b-d, SI Figure 3**). This is expected, as over an NK landscape, interactions between sites are not grounded in any real biophysical meaning and instead are made at random. We posit that only models with explicit structural understanding of the protein (be it learned in the model or through transfer learning) would be capable of meaningfully performing positional extrapolation.

### Sensitivity to sequence length

As a final test, the sensitivity of each model to sequence length was assessed. To test this, we performed a length expansion on sequences of length *N* from *NK* landscapes by adding consistent but arbitrary sequences in between *NK* sequence positions (**Figure 4a**). We tested sensitivity to sequence length for our chosen models against *NK* landscapes of increasing ruggedness with sequence lengths varying from 10 to 500 (**Figure 4b**). We see decision tree models being robust to sequence length, performing well up to *K* = 4 where even interpolation struggles (**Figure 2b-c**). Deep learning architectures are particularly affected, with MLPs and CNNs being the only models able to achieve reasonable predictive performance on the longest sequence lengths of 500 at *K* = 1, and, even so, exhibiting a decrease in performance compared to shorter sequence lengths of 10. The differences between decision tree models and deep learning models reflects fundamental architectural differences: RFs and GBTs create decision trees based on input features, identifying changing sequence positions as important and relegating the remainder of the sequence as inconsequential. The deep learning models show variation in performance with sequence length. For sequential models (LSTMs and Transformers) specifically, this is expected, as with increasing sequence length, interactions between sites become increasingly long-range and harder to model. However, even non-sequential neural network models experience a decrease in performance with sequence length increases. To assess if this trend was confounded by a decrease in performance due to the tuned hyperparameters becoming increasingly inappropriate for increased sequence lengths, we repeated the experiment on the CNN and bLSTM model on the *K* = 1 landscape, this time tuning the hyperparameters for the sequence length specifically. We see that the decrease in performance is mirrored even with hyperparameters tuned for the new sequence lengths, however, extreme dips in performance (for the bLSTM model) at sequence lengths of 200 and 250 are likely related to poor hyperparameter fitting (**SI Figure 5**). A decrease in performance at larger sequence lengths is not unsurprising. One possible reason for this is that an artificial increase in sequence length leads to additional parameters being added into the model that are not applied to meaningful variation in the data, instead capturing redundant features that cannot be used for generalized predictions. However, performance degradation due to an inflated feature space is also a likely contributing factor.^42,43^ It has been proven analytically that as the size of the sequence (i.e., the *N* parameter increases), so too does the ruggedness of the fitness landscape. For example, the number of expected local maxima in an NK landscape grows exponentially with *N* according to closed-form expressions,^44^ and the number and length of fitness-monotonic walks has been shown to grow logarithmically with *N*.^36^ However, we note that as we are not increasing the *N* parameter of the NK landscape, rather, arbitrarily padding the sequences to a given length, the ruggedness of the landscape is not affected, meaning decreasing performance is related to architectural challenges rather than the production of more complex landscapes.

**Figure 4.**
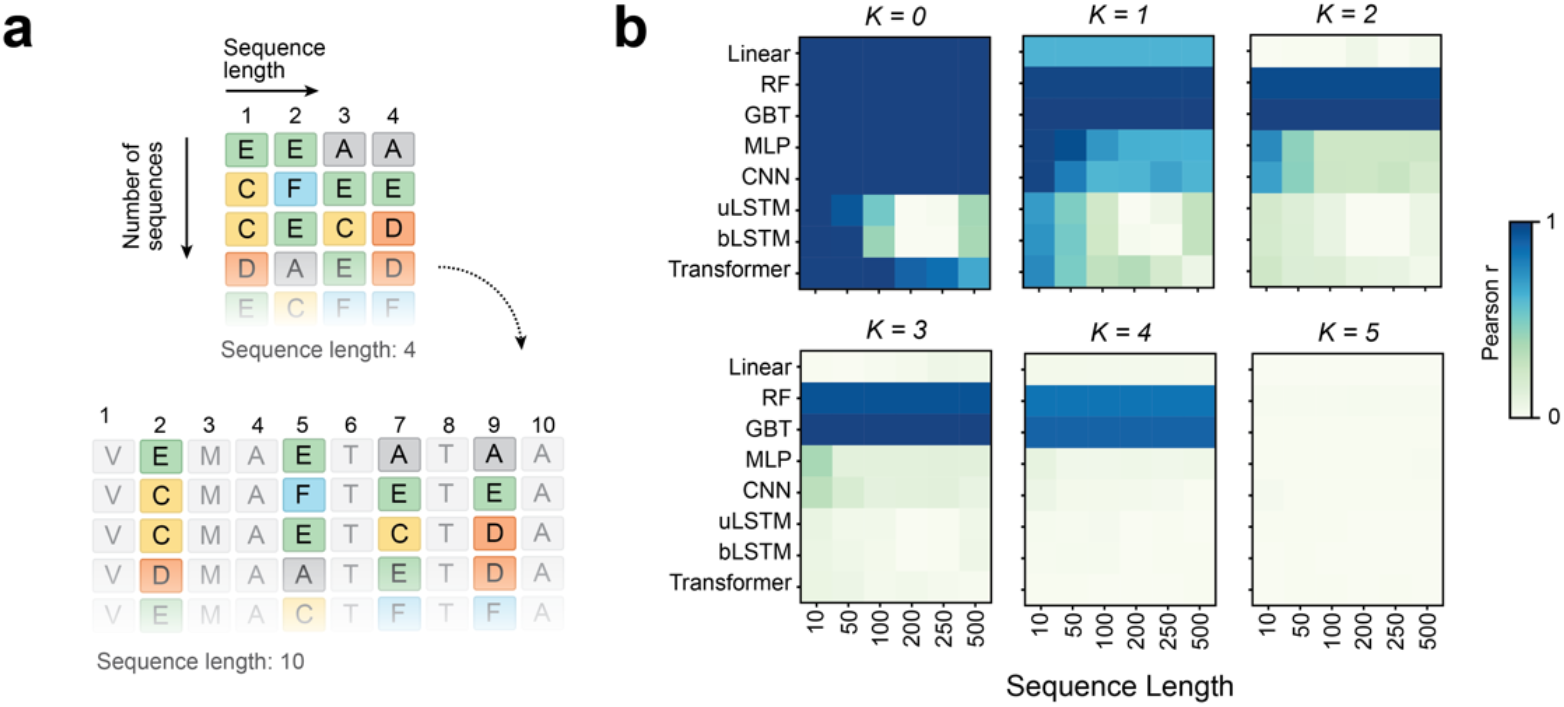
Sequence length dependency on NK Landscapes. (**a**) Schematic for sequence length adjustment. Briefly, arbitrary sequences are injected between positions to create a new sequence of a given length. While injected sequences are arbitrary, they are constant (i.e. unvarying) within a given landscape. (**b**) Sensitivity to sequence length for NK Landscapes with *K* between 0 and 5. Heatmaps show the model performance (Pearson *r*) on the test datasets for landscapes of increasing ruggedness (*K* = 0, . . ., 5).

### GB1 and extension to empirical datasets

Although the *NK* landscape is an excellent tool for evaluating the determinants ML sequence-fitness performance, its key limitation is that it is not rooted in biophysics. We therefore sought to validate results from the NK landscapes by comparing those results to an empirical protein fitness landscape. Generally, it is not experimentally tractable for any real protein dataset to exhaustively search three, let alone six mutational regimes. As a compromise, we used protein G domain B1 dataset (GB1) of Wu *et al*.,^45^ a combinatorial fitness landscape containing all 20 amino acids at 4 unique amino acid sites, thus spanning the sequence space (*M*_1_ − *M*_4_) at the 4 positions mutation (**Figure 5a**), permitting extrapolation to be tested on an experimental dataset. Indeed, the GB1 dataset has been used as a benchmark system for the analysis of epistasis and fitness landscape topography at length previously.^46–48^

When using the GB1 landscape, we can see that, similarly to NK landscape (**Figure 2c-d**), extrapolation performance (Pearson *r*) decreases as the distance of the extrapolation task extends (**Figure 5b-c, SI Figure 6**). This has important implications for training dataset structure in contexts where researchers are considering which sequences to assay in wet lab experiments in an ML pipeline. Here, sequences to assay should be those that are within the same mutational regimes as those desired to be predicted using ML, so that the ML model only must interpolate, rather than extrapolate.

**Figure 5.**
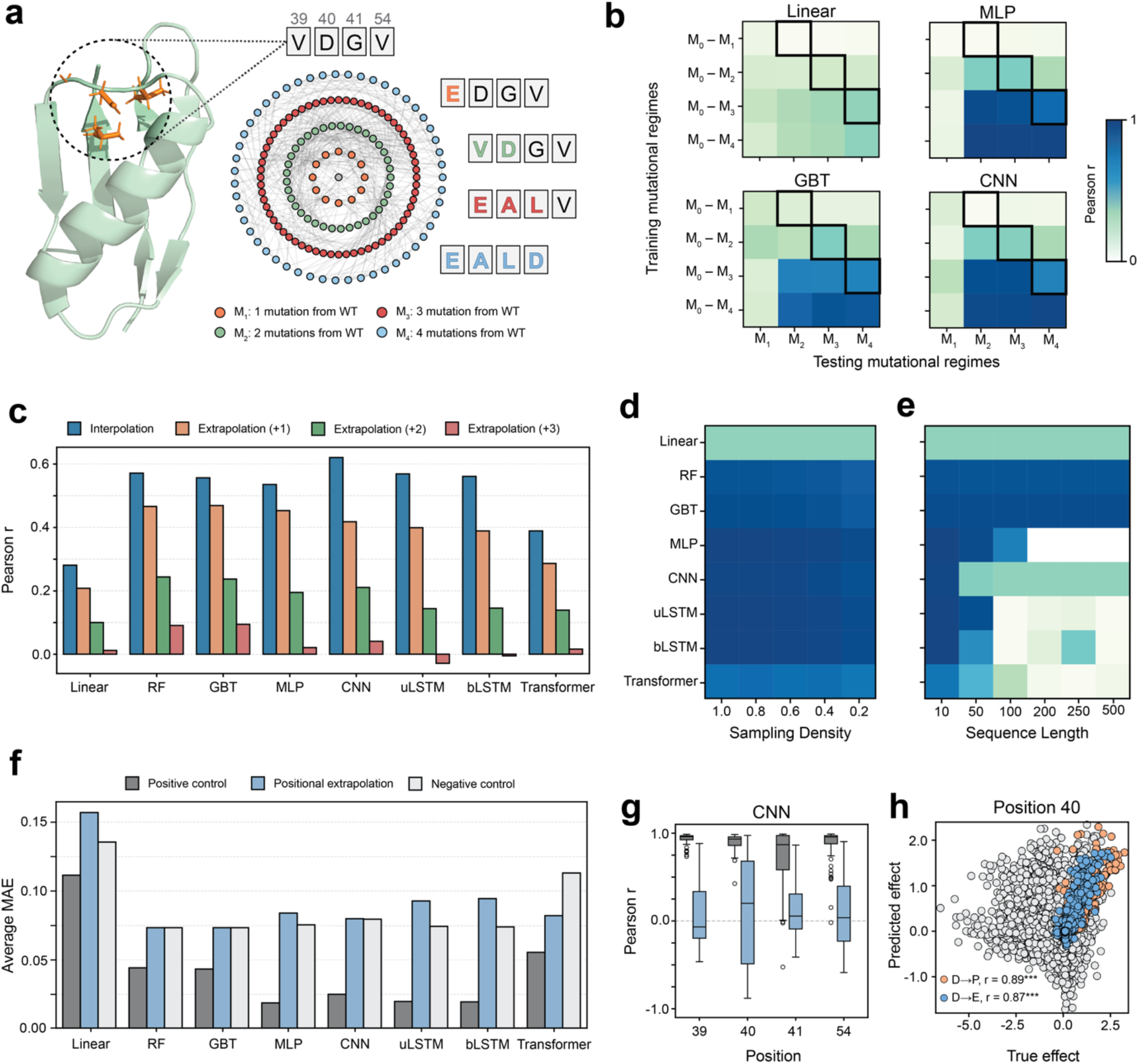
Interpolation and extrapolation on the GB1 landscape. (**a**) Solution NMR structure of *Streptococcus dysgalactiae* protein GB1 (PBD: 2GB1) showing mutated sites from the deep mutation scan dataset of *Wu et al* 2016 in orange, alongside a Hamming graph in radial layout showing mutational regimes from the central reference sequence (here, the wild-type sequence VDGV). (**b)** Heatmaps showing Linear, GBT, MLP and CNN model interpolation and extrapolation performance (Pearson *r*). Cells in a black border represent extrapolation into one mutational regime, to the left is interpolation and to the right extrapolation to higher mutational regimes. The y-axis of the heatmaps corresponds to training mutational regimes, and the x-axis to testing mutational regimes. (**c**) Bar plot showing interpolation and extrapolation to the first (+1), second (+2) and third (+3) mutational regimes beyond the training regimes, for all models. Performance is evaluated as the Pearson *r* on test data. (**d**) The performance (Pearson *r*) of all models as train data sampling density decreases, measuring robustness to sparse data. The y-axis of these heatmaps corresponds to the models tested, the y-axis to the test set sampling density. (**e**) Sensitivity to sequence length. Heatmap shows the model performance (Pearson *r*) on the test datasets. (**f**) The average MAE (mean absolute error) in predicting mutational effects (the fitness of the mutant minus the fitness of the WT) in the positional extrapolation testing regime for each model. (**g)** The correlation (Pearson *r*) between predicted mutational effects from the CNN model and true mutational effects per position. The negative control is not plotted as all correlation values are undefined (having predicted all mutational effects to be 0). (**h**) The predicted mutation effects from the CNN model against the true mutational effects at position 40 for one replicate (replicate 0, seed sequence DETN) separated for amino acid substitutions with the top two Pearson correlation values when comparing predicted and true mutational effects (for D40P in orange and D40E in blue) across all observed genetic contexts. Significance is annotated as: *** p < 0.001, ** p < 0.01, * p < 0.05.

As the percentage of landscape data used decreases, so too does the performance of all models (**Figure 5d**). Although this is to be expected, surprisingly, the extent of performance deterioration is minimal, especially for the neural network architectures (excepting transformers). Evidently, the GB1 landscape is sufficiently smooth to permit a small percentage of data to be highly informative to NN models regarding the overall structure of the landscape. In NK landscapes, we see that as landscape ruggedness increases, more data is needed and there is a sharper decay at lower sampling of the landscape. We see a similar trend in the GB1 landscape as we do in the NK landscape when increasing the sequence length (**Figure 5e**). Specifically, the resilience of decision tree models is maintained as well as the decrease in performance of neural networks as sequence length increases. We note that these results may differ if the full sequence of the GB1 protein were used (sequence length of 56 amino acids), where the information present may allow for improved learning of the protein-fitness landscape. For example, the additional sequence context provided by the full domain length may translate to more expressive representations of the individual mutations learned by deep neural networks (particularly by recurrence or attention mechanisms).

The GB1 landscape provides a unique opportunity to evaluate positional extrapolation in a combinatorial empirical landscape. Positional extrapolation is not possible on NK landscapes because interdependencies between sequence positions are not grounded in biophysics. In contrast, on a real protein landscape, information learned over one or more sites may be applicable to sites where no variation is present during training. When comparing the average MAE of the predicted mutational effects to the true mutational effects, all models have similar average values (**Figure 5f**) and error distributions (**SI Figure 7a**) to the negative control. However, when comparing the correlation (Pearson *r*) between predicted and true mutational effects at each site, some models appear to achieve some values of high correlation (**Figure 5g, SI Figure 7b**). Specifically, the CNN, MLP and transformer models, while having correlation values centralized on 0, had many instances of correlations greater than 0.8. We investigated this further on the CNN model (for which this observation was most pronounced) to determine if some level of positional extrapolation was occurring. When comparing the predicted effects to the true effects, strong significant correlations were observed for specific mutations (**Figure 5h, Figure 6c**). Predictions of this strength were not seen in the predictions of NK landscapes (**SI Figure 4**), and consequently, do not occur at random where no positional extrapolation is occurring. Interestingly, there are regular patterns across sites for which mutations were predicted well. Specifically, mutations to proline and/or glutamic acid displayed strong correlations, indicating successful positional extrapolation. (**Figure 5h, SI Figure 7c**). To map this out further, we repeated the positional extrapolation experimentation on the CNN model with each amino acid fixed at each site. The results of these experiments show that the CNN model has a bias in being better able to predict positive fitness effects (**SI Figure 8, 9**).

Earlier, we proposed that models likely need to learn structural and biophysical features of a given protein system to achieve performance on positional extrapolation tasks. To assess this, we compared the representations of individual amino acids in the latent space of the CNN with one-hot embeddings (OHE) (**SI Figure 10**). Our results indicate that the CNN latent space is able to cluster amino acids based on biophysical properties and size, a well-known phenomenon in protein representation learning.^3^ While the NK landscape cannot be used to benchmark models on positional extrapolation tasks (given its lack of biophysical grounding), it is useful in serving as a negative (random) control for distinguishing random successes from true predictive capabilities for models predicting on empirical fitness landscapes. By comparing the positional extrapolation results on NK landscapes (**Figure 3**) with the GB1 positional extrapolation results, we can conclude that some models are able to perform genuine positional extrapolation for some positional mutations on empirical landscapes.

## Discussion

### The central role of ruggedness

From our results, ruggedness emerges as the key determinant of ML model performance, with greater effect on performance than any other tested variable. This is not surprising: learning depends on extracting regularities in data and using these regularities to inform predictions. Indeed, previous studies have commented on the decay of model performance as a function of landscape ruggedness.^49^ As epistasis increases, the fitness function becomes more uncorrelated, with individual mutations having outsized and unpredictable effects on fitness. Fundamentally, all ML sequence-fitness predictors approximate the fitness function; when the landscape is maximally rugged, the NK model assigns fitnesses at random such that covariation between sequences does not exceed the sampling variance (i.e. it is equivalent to sampling randomly from some fitness distribution). Indeed, when *K* = *N* − 1, the NK landscape reduces to a “house-of-cards” model of epistasis, where fitness labels are independent and identically distributed samples drawn at random from an underlying fitness distribution and therefore share no covariation.^44^ Generally, for supervised ML methods (including the models used here), as the complexity/irregularity of the function being approximated increases, the accuracy of approximation deteriorates if the number of training samples and model capacity (e.g. number of neurons, number of layers) are kept constant.^50–53^ More training samples are therefore needed (along with greater model capacity) for a model to effectively learn complex functions. This has been described previously in quantifying the ruggedness of fitness landscapes^7,54^. Indeed, in the limit of a completely uncorrelated and random function, the number of training samples required to accurately predict approaches the size of the domain on which prediction is being performed. This is a key result of Kolmogorov complexity: a completely random function has no shorter description than an explicit list of its values.^50^

A corollary of this is that, in the case of proteins, as epistasis becomes more pervasive (and thus the fitness function’s complexity increases), more training samples are needed to sustain performance. Our results in **Figure 2d** are consistent with this fact. This fundamental dependence of ML performance on landscape ruggedness highlights the need for a clear understanding of ruggedness in experimental landscapes. If the ruggedness of a given experimental landscape can be characterized, a clearer prediction of ML performance on that landscape can be made. Several strategies exist for calculating the ruggedness of protein fitness landscapes, including number of local maxima,^55,56^ graph Dirichlet energy,^57,58^ Fourier transform,^59^ and *r/s* ratio,^60^ among others.^19,27,61–67^ For most of such approaches, combinatorial completeness (or near completeness) is often a prerequisite, hindering practicality. Graph Dirichlet energy stands out as an approach that can effectively measure ruggedness without requiring combinatorial completeness,^19,29^ offering a simple approach for workers in the field wishing to calculate the ruggedness of their datasets prior to using them in an ML sequence-fitness pipeline. We note, however, that the Dirichlet energy varies with fitness scale and its interpretation is influenced by the underlying graph connectivity scheme (i.e. the graph topology).

### Recommendations when designing experimental training data

We can make a few recommendations for designs of wet-lab experiments as part of ML-driven sequence-fitness prediction pipelines. The first recommendation is consideration of mutational regime (MR) extrapolation and interpolation. When designing experiments, training data should sample the mutational regimes that the ML model will be foreseeably asked to predict within. That is, experimental sampling should be designed in such a way as to avoid asking the ML models to extrapolate. If extrapolation is unavoidable, care should be taken to minimize its extent (i.e. only to extrapolation +1, +2 etc.). In this way, ML performance can be maximized.

Secondly, the extent of sampling (i.e. sampling density) should be considered in the context of the degree of epistasis/ruggedness present in the experimental protein fitness landscape. Where ruggedness is high, greater sampling will be required. Nevertheless, if the protein GB1 dataset is representative of empirical datasets in general, our results on this empirical dataset suggest that sampling a relatively small proportion of the fitness landscape (~20%) can yield excellent results. Further research on other empirical datasets is needed to establish a clearer and more general impression regarding the degree of sampling required to obtain good results; nevertheless, existing literature does offer insight on the challenges of sparse data sampling and the interpretations of epistasis and ruggedness that can be gleaned from incomplete data.^48,54,67^

Interestingly, our results show that positional extrapolation is possible using NN architectures on experimental landscapes (**SI Figure 7b**). This is possible because these NN models can implicitly learn biophysical properties of amino acids seen at other sites (**SI Figure 10)** and extrapolate these properties to sites where they have not seen the amino acid(s) in question. Nevertheless, when designing experiments, training data should sample a diverse array of amino acid categories (e.g. hydrophobic, polar, aromatic etc.) at each site.

When considering the capabilities of the CNN model in predicting mutational effects, a bias towards success in predicting positive mutational effects was observed (**SI Figure 8, 9**). This bias may be related to the use of Mean Square Error (MSE) loss function biasing the model to achieve higher performance when predicting high fitness values. Consideration of the impacts of loss functions as well as data label skew may be necessary to assess if such biases may be introduced.

In this work, when performing positional extrapolation, we held the identity of the amino acid at the *i*^*th*^ site constant during training, and tasked models with predicting the mutational effect of alternative amino acids at that *i*^*th*^ site; an interesting future experiment could be to sample single representative amino acids from each category at the *i*^*th*^ site and then to task the model with predicting the mutational effect of alternative amino acids at that *i*^*th*^ site. This strategy would provide general information to ML models regarding how categories of amino acids at the *i*^*th*^ site affect fitness, theoretically permitting models that have implicitly learnt biophysical properties of amino acids to perform better at positional extrapolation. If there is a meaningful improvement in performance from such an experiment, a recommendation could be made to sample at least one amino acid from each biophysical category at each position when designing experiments.

A key limitation of the *NK* landscape here is its lack of biophysical grounding, meaning that positional extrapolation is not possible on *NK* landscapes. Future work on identifying/developing model fitness landscapes with grounding in biophysics (e.g. generalized/structurally informed NK^7^ and NKp^40^ models) could be useful as alternatives to the *NK* landscape when assessing the ability of NN models to perform positional extrapolation.

Finally, models vary in their suitability for a particular goal (i.e. interpolation vs extrapolation vs robustness to sparse training data vs positional extrapolation). We find that decision trees have the best performance on interpolation and extrapolation tasks and are most robust to sparse training data but are not able to perform positional extrapolation (which NN models excel at). Of the NN models, MLP and CNN are the best all-rounder models. Another important consideration for modelling is computational efficiency and wall-time. We found that the GBT and Transformer models had the greatest wall-time, followed by the LSTM models. It is well-known that GBT models become increasingly inefficient with larger datasets; close alternatives such as histogram-based gradient boosting regressors or parallelized strategies such XGBoost can be used to overcome this limitation. In terms of pure computational efficiency on larger datasets, random forests, MLP and CNN excel.

### Protein language models, regression models and ruggedness of representations

Increasingly, the field has embraced protein language models (PLMs) such as UniRep,^17^ ESM,^3,5,68^ LASE,^49^ and RELSO^69^ to represent proteins in semantically rich embedding spaces prior to using top-model regressors for sequence-fitness prediction tasks. While outside the scope of this work, we hypothesize that using PLMs will permit better positional and MR extrapolation due to the input spaces’ implicit encoding of biophysical and evolutionary properties.^70^ Future work on characterizing the performance boost (if any) rendered by PLMs on regressors performing positional and MR extrapolation will be deeply interesting and useful to the field.

Finally, preliminary data indicates that PLMs can generate smooth representations of protein fitness landscapes in their embedding spaces compared to the ruggedness of the input protein fitness landscapes,^49,69^ and that the extent of this smoothening correlates with performance.^49^ Indeed, strategies that minimize ruggedness of embedding spaces explicitly are well-known in machine learning for producing good results.^7,67,69^ It has been demonstrated that deep learning models like AlphaFold2^1^ and RoseTTAFold^71^ can improve protein binder design success rates nearly 10-fold by assessing the probability that designed sequences adopt the intended structure and binding mode. This suggests that these models effectively smooth the fitness landscape by capturing underlying patterns that physical energy-based methods may miss.^71^ It is tempting to speculate that, to the extent a PLM (or indeed a regressor model) generates a smooth representation of a protein fitness landscape in its embedding space, it will perform better on both positional and MR extrapolation. Future work on this issue will be illuminating and useful to the field.

## Conclusion

In conclusion, this work has outlined a principled approach to experimental data sampling, model selection and evaluation rooted in fitness landscape theory. By measuring model performance on synthetic (*NK*) and empirical fitness landscapes along the metrics of interpolation, MR extrapolation, positional extrapolation and sensitivity to sparse training data, we have shown that ruggedness emerges as the preeminent determinant of model performance, consistent with the tenets of approximation theory. Based on our results, we have outlined recommendations for designing experimental data, namely that (1) training data should be designed to minimize extrapolation, (2) the required training data sampling density depends on the ruggedness of the fitness landscapes, and that for empirical datasets, good performance can be attained with low (~20%) sampling, (3) although positional extrapolation is possible with NN architectures (but not decision-tree models), diverse sampling of amino acid categories at each position will likely result in improved performance, and (4) RF, MLP and CNN models are excellent initial choices that maximize performance without the computational cost/wall-time performance issues of alternative models. Finally, we have outlined promising areas of future work, including establishing the possible performance boost conferred by PLM models along the metrics outlined in this work. We hope the findings of our work will provide the field with a set of strategies and metrics that help meaningfully advance ML sequence-fitness prediction accuracy, with valuable applications in protein engineering and medicine.

## Methods

### The NK Landscape

The NK landscape of Kauffman and Weinberger^33^ is a theoretical model of combinatorial fitness landscapes that captures the key characteristic of these landscapes: *ruggedness*, which results from *epistasis. N* gives the number of positions in the sequence (i.e. length of the sequence); *K* gives the number of positions that interact with a given position *i*; thus, *K* gives the order of epistasis in the landscape. For any position *i*, the set of positions that interact with it are *ν* = {*ν*_*i*,1_, *ν*_*i*,2_, . . ., *ν*_,*Y*_}. These are typically assigned uniformly and independently at random. The fitness of some sequence X of length *N* is given by Equation 1:

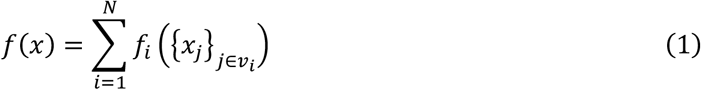

Here 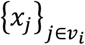 is the set of positions that position *i* interacts with (i.e. the interaction set). If any of the positions in the interaction set for some position *i* are mutated, a new fitness *f*_*i*_ is randomly assigned from a continuous uniform distribution on the interval [0, 1); else, *f*_*i*_ stays the same. It then follows that as the interaction set of *i* increases (with increasing *K*), the probability that a mutation anywhere in the sequence will affect the fitness contribution *f*_*i*_ of position *i* also increases (i.e. the likelihood of *f*_*i*_ being re-assigned increases). This means that a single mutation affects the sequence fitness *f*(*X*) more drastically, thereby increasing the ruggedness of the fitness landscape.

All NK landscapes were generated using custom code that was adapted from Obolski and colleagues.^72^ For each ruggedness value, eight landscapes were generated from randomized initial conditions. Each of these landscapes contained all sequences of length six (*N* = 6), using the first 6 canonical amino acids of the single letter amino acid alphabet. Limiting the amino acid pool was necessary to restrict the population size to a number that is tractable for many ML architectures, while also maintaining a sequence length that enables a wide variety of ruggedness values.

### Experimental GB1 dataset

To validate outcomes over the NK landscape for ML sequence-fitness prediction performance, we compared ML model performance qualitative rankings the B1 domain of immunoglobin-G (IgG) binding protein G (hereafter referred to as protein GB1),^45^ an empirical dataset from deep sequencing. Protein GB1 is a well-characterized prokaryotic protein extensively used as a model system in protein science, along with use in biotechnology applications like antibody purification. We used the deep mutational scan dataset from Wu and colleagues that exhaustively samples the sequence space of 4 positions (positions V39, D40, G41 and V54) near the C-terminus of the domain, consisting of 149361 (**SI Table 1**) variants. Fitness was calculated according to equation 2. In our framework, this dataset spans the first 4 mutational regimes (*M*_1_ − *M*_4_) in the sequence space of these 4 positions (i.e. it spans the entire sequence space of these 4 positions). We refer to these datasets as protein GB1. Both the input and selection counts were transformed into fitness values using Equation 2, with input and selection count being used as defined by.

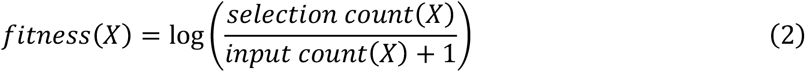

### ML model architectures

To demonstrate broad validity of this approach, we tested the NK landscapes against a wide variety of established ML architectures. The models fell broadly into three categories: (1) the decision-tree based architectures random forest (RF)^73^ and gradient boosted trees decision tree regression (GBT)^74^; (2) deep learning architectures including a multilayer perceptron (MLP)^75^, convolutional neural networks (CNN)^76,77^, unidirectional and bidirectional long short-term memory neural networks (uLSTM and bLSTM respectively)^78^, and transformer models^79^; and (3) a linear regression method, referred to simply as Linear. All models were implemented in scikit-learn (RF and GBTs) or PyTorch (all remaining models). All models were made without encoding layers, instead receiving one-hot embeddings (OHE) of sequence data to ensure comparisons being made were not biased by embedding layers being amenable to some models and not others.

### Model hyperparameter tuning

The hyperparameters of each model were tuned for each landscape. For NK landscapes, tuning was applied to a single replicate (with a 20% holdout validation set) and then used for training later models with the same *K* value. All models were tuned with Optuna 4.0.0. Specifically, hyperparameters were tuned with the Tree-structured Parzen Estimator (TPE) algorithm. The optimization process was initialized with 10 random trials, after which 15 × 2 (for decision tree models) or 15 × # *hyperparameters* (for neural networks) additional trials were conducted using the TPE approach. The hyperparameter search space for each model and landscape is specified in **SI Table 2**.

### Interpolation and extrapolation

Both interpolation and extrapolation were tested using the same testing regime. Mutational regimes were defined as being groups with sequences that shared the number of mutations from a seed sequence (for instance, *M*_1_ contained all variants with 1 mutation on the seed sequence). For each regime, 20% of sequences were held out as testing data and the remainder were used for training. Then, training and testing was conducted by accumulating the train data from *n* regimes (for example, for *n* = 3, training data from *M*_0_, *M*_1_, *M*_2_, and *M*_0_ would make up the training data) for testing on the test data of each regime separately. This testing regime was run with 4 replicates per landscape, where the seed sequence varied in each replicate to produce a new set of mutational regimes.

### Sensitivity to ablation

Ablation testing sought to determine the performance of each model on increasingly sparse data over a range of density values (100%, 80%, 60% 40% and 20% of the original data). After producing a 20% hold out test dataset, the density of the data was reduced by randomly extracting a percent of the full train dataset. Testing was then run over the full test split. For each landscape, 4 replicates were run where the seed sequence varied to produce a new set of mutational regimes.

### Positional extrapolation

We define positional extrapolation as the ability of a model to predict the influence of a mutation at a position that was not altered in the training set. To test this, we compared the true effect of a mutation to the predicted effect (Equation 3):

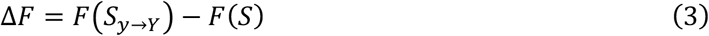

Where Δ*F* is the change in fitness (mutational effect), *F*(*S*_*y*→*Y*_) is the fitness of the mutant where the amino acid at position *i* has been modified, and *F*(*S*) is the fitness of the seed sequence with the original amino acid at position *i*. When testing positional extrapolation performance across models, for each landscape, 4 replicates were run, resulting in 4 seed sequences per run (and therefore the effects of mutations upon up to 4 amino acids per site being tested).

Models were trained for each site, where sequences with the amino acid of the seed sequence fixed at the site being set as training data. The model was then tested on each possible mutation at that site, measuring the mean error of prediction and the Pearson *r* to evaluate success. To produce a positive control, the same training and testing regime was repeated; however, 80% of sequences containing a non-seed amino acid at the test site were removed from the testing data and incorporated into the training data. This was done over each possible mutation at the site such that there were equal amounts of each mutant amino acid in the training data (and equal test sizes for each mutation). To produce a negative control, decision tree models were trained on shuffled data, and neural networks were not trained.

To determine the performance of the CNN model in predicting all mutations at each site of the GB1 protein, positional extrapolation was repeated with seed sequences that spanned all amino acids at all sites (*AAAA, CCCC*, …, *YYYY*).

### Sensitivity to sequence length

To test sequence length sensitivity, sequence lengths were artificially increased by randomly injecting NK or GB1 data into randomly produced sequences of a given length. Here, data injections were into arbitrary but constant positions within a fixed sequence. While the NK and GB1 landscape had sequence lengths of 6 and 4 respectively, they were both tested on sequence lengths of 10, 50, 100, 200, 250, and 500. Testing was conducted with a 20% holdout set with 4 replicates, where the seed sequence varied to produce a new set of mutational regimes.

### Landscape and amino acid visualization

Both full combinatorial landscapes and amino acids were represented in 2-dimensions with Uniform Manifold Approximation and Projection (UMAP). For combinatorial landscapes, OHEs of full sequences were reduced with UMAP. To embed amino acids in the latent space of the model trained on full amino acid sequences, OHE were altered to contain a vector containing the amino acid of interest at site one, and zero vectors at all other sites, and fed into the model as inputs. To embed amino acids in OHE, single vectors containing each amino acid individually were used as input to UMAP. UMAP was conducted with

### Statistical tests

The equality between MAE distributions between positional extrapolation and the positional extrapolation positive and negative control were tested using the Kolmogorov-Smirnov test, where a value of 1 indicates the two distributions are from different underlying distributions and 0 indicates they are from the same. Correlation was measured with Pearson’s correlation coefficient. Both tests were conducted with SciPy 1.14.1.

## Code Availability

All code used for data simulation, model training, hyperparameter tuning, and model benchmarking is available on GitHub (https://github.com/RSCJacksonLab/nk-ml).

## Acknowledgements

A.C.M. thanks the Australian National University and the Westpac Scholars Trust for PhD scholarships. This project was funded by the RAC Centre of Excellence in Peptide and Protein Science. The funders had no role in study design, data collection and interpretation of the decision to submit the work for publication.

## Author Contributions

A.C.M., M.S. and C.J.J. conceived the study. A.C.M., M.S. and D.S.M. conceived the experiments and methods, wrote the software, analyzed results, prepared figures and wrote and edited the manuscript. A.C.M. unified and vectorized the protein landscape class. A.C.M., M.S., and D.S.M., acquired and applied computational resources and performed the simulation experiments. M.A.S., A.A.L., and C.J.J. reviewed and edited the manuscript. C.J.J. supervised and administered the project and acquired funding.

