## Supplementary Information for "Investigating the determinants of performance in machine learning for protein fitness prediction"

**SI Table 1. Dataset properties.**

| Dataset | Number of datapoints | Alphabet size | Sequence length |
|---|---|---|---|
| **NK Landscapes (K = 0-5)** | **46 656** | **6** | **6** |
| Mutational regime 0 | 1 | | |
| Mutational regime 1 | 30 | | |
| Mutational regime 2 | 375 | | |
| Mutational regime 3 | 2 500 | | |
| Mutational regime 4 | 9 375 | | |
| Mutational regime 5 | 18 750 | | |
| Mutational regime 6 | 15 625 | | |
| **G Domain B1 dataset (GB1)** | **149 361** | **20** | **4** |
| *Theoretical dataset size | 160 000 | | |
| Mutational regime 0 | 1 | | |
| Mutational regime 1 | 76 | | |
| Mutational regime 2 | 2 166 | | |
| Mutational regime 3 | 27 436 | | |
| Mutational regime 4 | 130 321 | | |

The number of datapoints, alphabet size and sequence length for the NK Landscape and GB1 datasets. The number of datapoint sizes present in mutational regimes (referred to as $M_0 - M_6$) throughout are reported. Noting that the mutational regime sizes reported for the GB1 B1 dataset are calculated on the theoretical dataset size, as the empirical dataset has missing values.

**SI Table 2. Hyperparameter search space for model and dataset**

| Dataset | Model | Search space |
|---|---|---|
| NK Landscapes (K= 0-5) | Random Forest | Number of trees: [10, 1000] (integer range, inclusive)<br>Maximum fraction of features: [0.1, 1] (float range, inclusive)<br>Maximum tree depth: [1, 32] (integer range, inclusive) |
| | Gradient Boosted Trees | Number of trees: [10, 1000] (integer range, inclusive)<br>Maximum tree depth: [1, 32] (integer range, inclusive)<br>Learning rate: [0.001, 0.2] (float range, inclusive) |
| | Linear | Learning rate: {0.01, 0.001, 0.0001}<br>Batch size: {32, 64, 128, 256} |
| | MLP | Learning rate: {0.01, 0.001, 0.0001}<br>Batch size: {32, 64, 128, 256}<br>Number of hidden layers: [1, 3] (integer range, inclusive)<br>Hidden layer size: {32, 64, 128, 256} |
| | CNN | Learning rate: {0.01, 0.001, 0.0001}<br>Batch size: {32, 64, 128, 256}<br>Number of convolutional layers: [1, 2] (integer range, inclusive)<br>Number of kernels per layer: [32, 256] (integer range, increments of 32, inclusive)<br>Kernel size: [3, 5] (integer range, inclusive) |

| | | |
|---|---|---|
| | LSTM (unidirectional and bidirectional) | Learning rate: {0.01, 0.001, 0.0001}<br>Batch size: {32, 64, 128, 256}<br>Number of layers: [1, 2] (integer range, inclusive)<br>Hidden units per layer: {64, 128, 256} |
| | Transformer | Learning rate: {0.01, 0.001, 0.0001}<br>Batch size: {32, 64, 128, 256}<br>Embedding dimension: {32, 64, 128, 256}<br>Number of attention heads: [1, 8] (integer range, inclusive)<br>Number of layers: [1, 2] (integer range, inclusive)<br>Hidden layer dimension: {32, 64, 128, 256} |
| G Domain B1 dataset (GB1) | Random Forest | Number of trees: [10, 1000] (integer range, inclusive)<br>Maximum fraction of features: [0.1, 1] (float range, inclusive)<br>Maximum tree depth: [1, 32] (integer range, inclusive) |
| | Gradient Boosted Trees | Number of trees: [10, 1000] (integer range, inclusive)<br>Maximum tree depth: [1, 32] (integer range, inclusive)<br>Learning rate: [0.001, 0.2] (float range, inclusive) |
| | Linear | Learning rate: {0.01, 0.001, 0.0001}<br>Batch size: {32, 64, 128, 256} |
| | MLP | Learning rate: {0.01, 0.001, 0.0001}<br>Batch size: {32, 64, 128, 256}<br>Number of hidden layers: [1, 6] (integer range, inclusive)<br>Hidden layer size: {32, 64, 128, 256, 512} |
| | CNN | Learning rate: {0.01, 0.001, 0.0001}<br>Batch size: {32, 64, 128, 256}<br>Number of convolutional layers: [1, 2] (integer range, inclusive)<br>Number of kernels per layer: [32, 512] (integer range, increments of 32, inclusive)<br>Kernel size: {3} |
| | LSTM (unidirectional and bidirectional) | Learning rate: {0.01, 0.001, 0.0001}<br>Batch size: {32, 64, 128, 256}<br>Number of layers: [1, 2] (integer range, inclusive)<br>Hidden units per layer: {64, 128, 256, 512} |
| | Transformer | Learning rate: {0.01, 0.001, 0.0001}<br>Batch size: {32, 64, 128, 256}<br>Embedding dimension: {32, 64, 128, 256, 512}<br>Number of attention heads: [1, 8] (integer range, inclusive)<br>Number of layers: [1, 3] (integer range, inclusive)<br>Hidden layer dimension: {32, 64, 128, 256, 512} |

Model hyperparameters were tuned with the Tree-structured Parzen Estimator (TPE) algorithm. The optimization process was initialized with 10 random trials, after which 15 × 2 (for decision tree models) or 15 × number of hyperparameters (for neural networks) additional trials were conducted using the TPE approach. Hyperparameter performance was assessed against 20% of the training data as a validation set.

**SI Table 3. Number of model parameters.**

| Landscape | Model | Number parameters |
|---|---|---|
| NK Landscape, K=0 | Linear | 37 |
| | MLP | 2 433 |
| | CNN | 208 353 |
| | uLSTM | 69 761 |
| | bLSTM | 541 185 |
| | Transformer | 265 985 |
| NK Landscape, K=1 | Linear | 37 |
| | MLP | 70 785 |
| | CNN | 180 417 |
| | uLSTM | 270 593 |
| | bLSTM | 534 785 |
| | Transformer | 42 305 |
| NK Landscape, K=2 | Linear | 37 |
| | MLP | 17 729 |
| | CNN | 291 489 |
| | uLSTM | 270 593 |
| | bLSTM | 541 185 |
| | Transformer | 100 481 |
| NK Landscape, K=3 | Linear | 37 |
| | MLP | 42 497 |
| | CNN | 180 417 |
| | uLSTM | 201 857 |
| | bLSTM | 541 185 |
| | Transformer | 265 985 |
| NK Landscape, K=4 | Linear | 37 |
| | MLP | 15 105 |
| | CNN | 255 585 |
| | uLSTM | 796 929 |
| | bLSTM | 2 118 145 |
| | Transformer | 42 305 |
| NK Landscape, K=5 | Linear | 37 |
| | MLP | 4 481 |
| | CNN | 30 817 |
| | uLSTM | 18 497 |
| | bLSTM | 2 118 145 |
| | Transformer | 100 481 |
| G Domain B1 dataset (GB1) | Linear | 81 |
| | MLP | 435 713 |
| | CNN | 350 113 |
| | uLSTM | 1 094 145 |
| | bLSTM | 2 188 289 |
| | Transformer | 267 777 |

The number of parameters for hyperparameter tuned neural network architectures for each landscape.

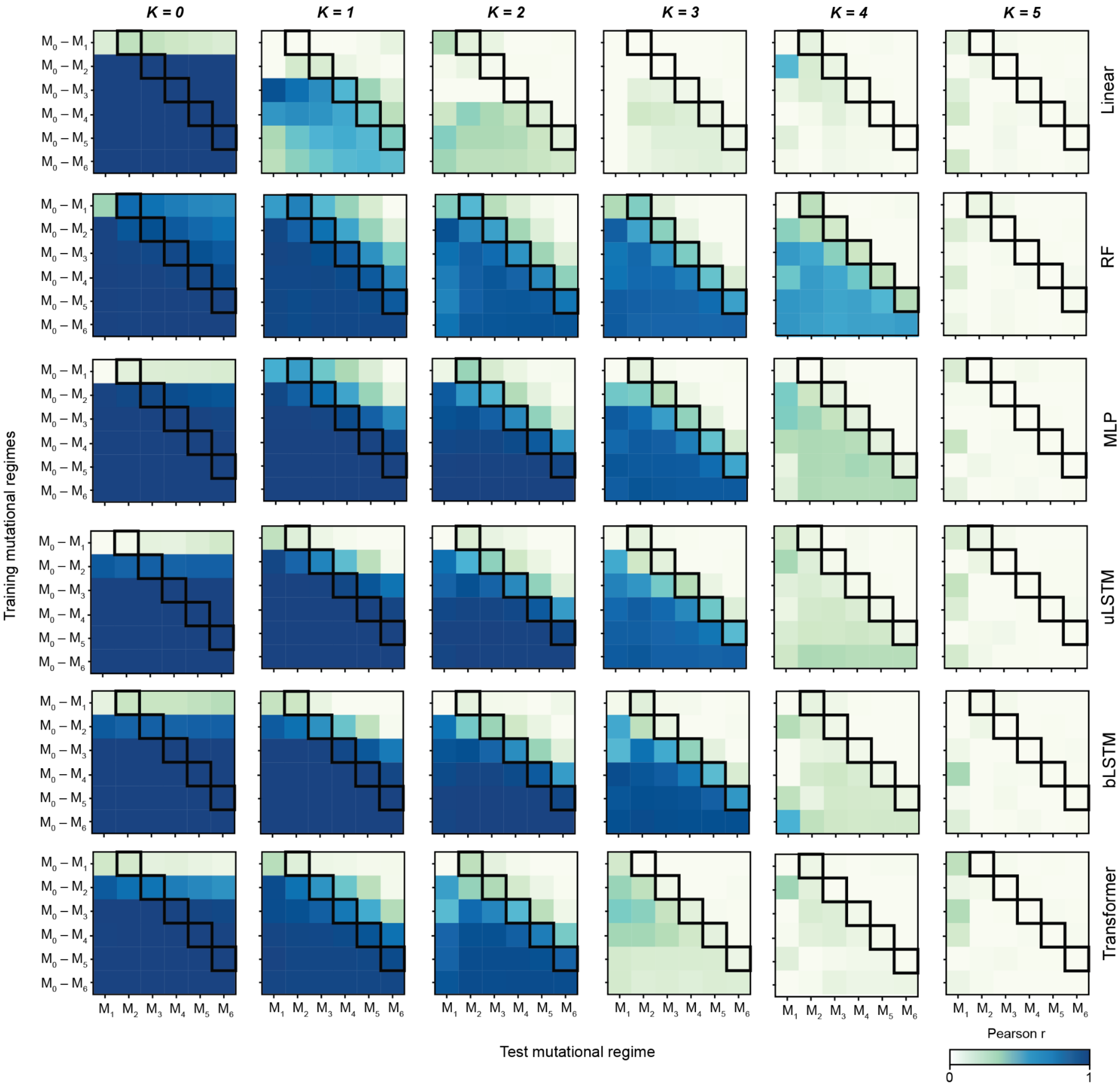


**SI Figure 1. Interpolation and extrapolation over the NK Landscapes.** The average Pearson's correlation (over 4 replicate NK landscapes) for the GBT and CNN models. Cells in a black border represent extrapolation into one mutational regime (variants containing one additional mutation compared to the training data), to the left is interpolation and to the right extrapolation to higher mutational regimes (variants containing two to five additional mutation compared to the training data). The y-axis of the heatmaps corresponds to training mutational regimes, and the x-axis to testing mutational regimes.

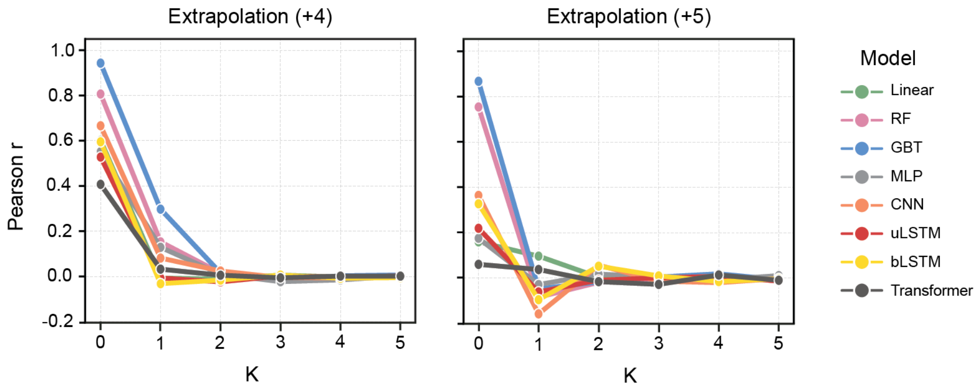


**SI Figure 2. Extrapolation to +4 and +5 mutational regimes on the NK Landscape.** The performance (Pearson *r*) of models over increasing ruggedness (K) on extrapolation +4 and extrapolation +5 (variants containing four or five additional mutations compared to the training data respectively) test datasets.

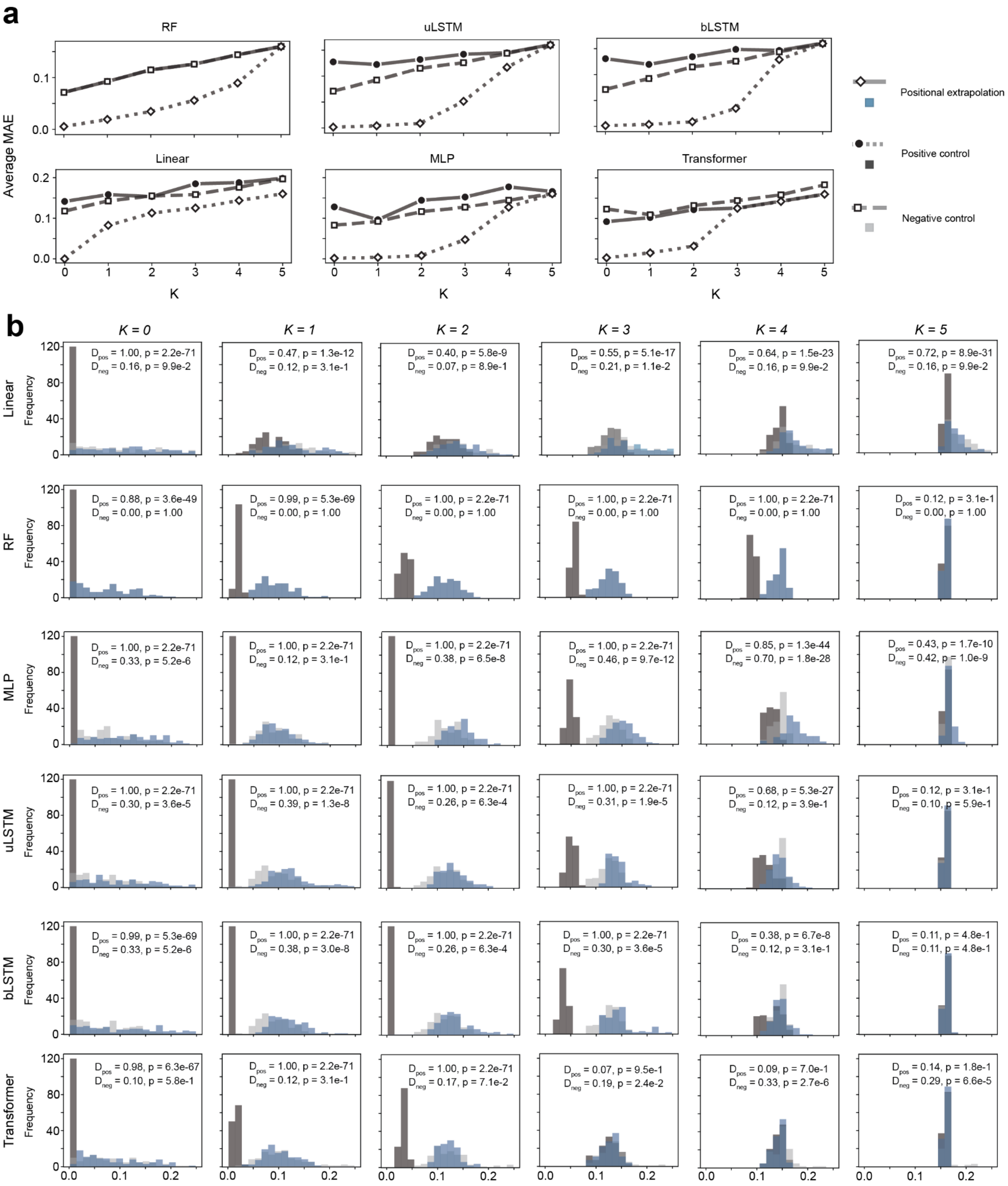


**SI Figure 3. Positional extrapolation performance (MAEs) over the NK landscape** (**a**) The average MAE (mean absolute error) of linear, RF, MLP, uLSTM, bLSTM and transformer models in predicting mutational effects (the fitness of the mutant minus the fitness of the WT) in the positional extrapolation testing regime against K. The negative control overlays the positional extrapolation data for the GBT model. (**b**) Histogram of MAEs for linear, RF, MLP, uLSTM, bLSTM and transformer models when predicting mutational effects in the positional extrapolation testing regime. D is Kolmogorov's D statistic when comparing the MAEs associated with positional extrapolation against the negative ($D_{neg}$) and positive ($D_{pos}$) control. D is 1 for sample distributions drawn from separate underlying distributions and 0 for those drawn from the same distribution. P-values < 0.05 are considered to be statistically significant. The positive control represents data with the same splitting regime with 80% of sequences containing amino acids other than the fixed amino acid reintroduced to the training dataset and the remaining 20% of sequences used as a test dataset. The negative control represents neural network models with randomly initialized weights or decision tree models trained with shuffled training targets.

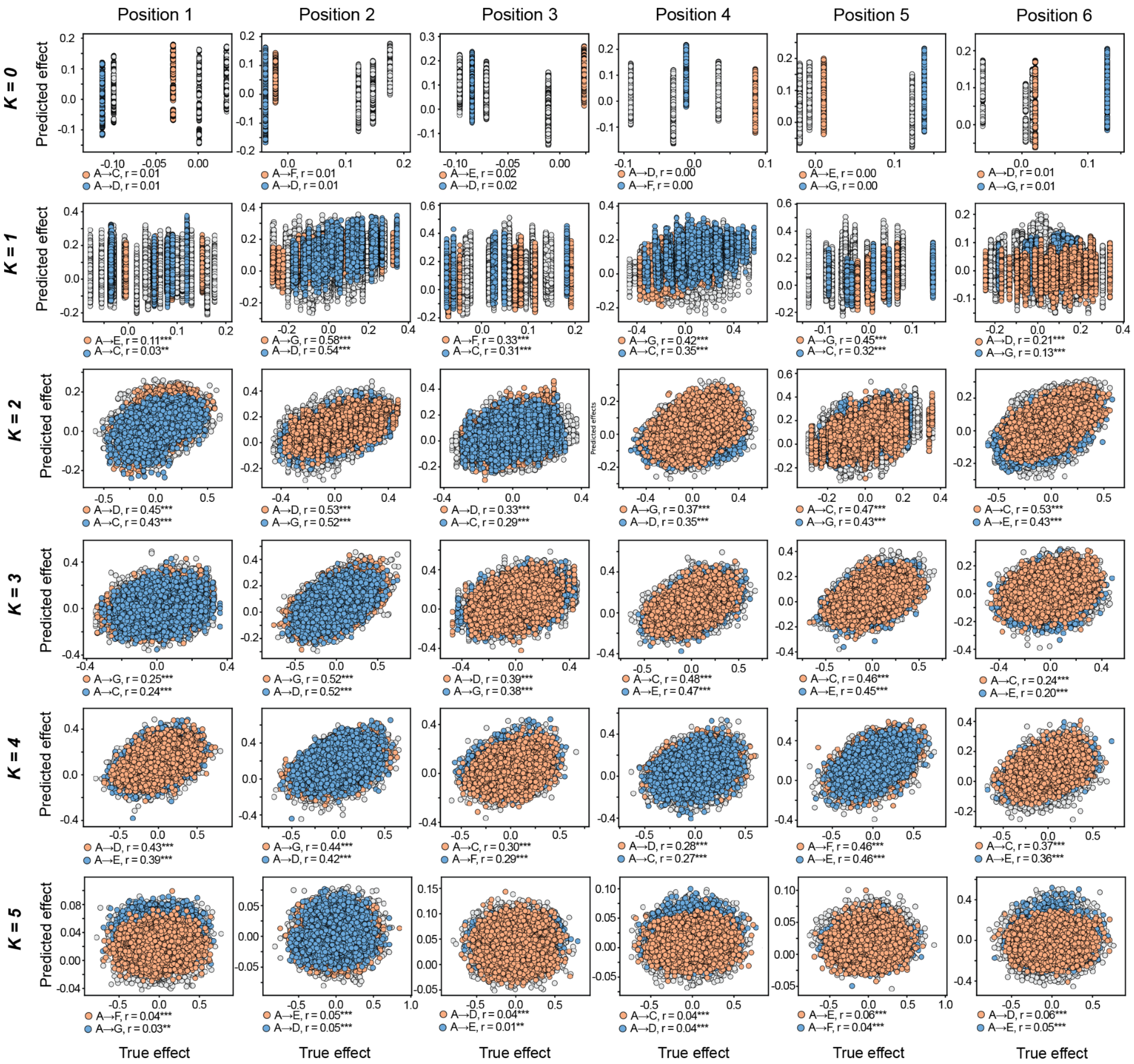


**SI Figure 4. Predicted mutational effects by CNN models on the NK Landscape.** The predicted mutation effects from the CNN model against the true mutational effects at each position (1-6) in NK simulated datasets. Positional extrapolation is separated for amino acid substitutions where the subsequent effect was predicted resulting in a correlation greater than 0.8 across all observed genetic contexts. Significance is annotated as: *** $p < 0.001$, ** $p < 0.01$, * $p < 0.05$.

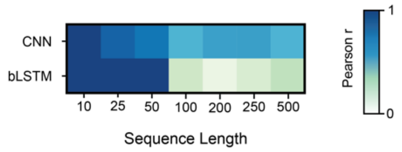


**SI Figure 5. Sequence length dependency of CNN and bLSTM models with hyperparameter tuning.** Heatmap showing the performance (Pearson *r* on test data) of CNN and bLSTM models as the sequence length of the data increases on the K=1 landscape. For each sequence length, model hyperparameters were tuned on the replicate 0 landscape and the same 4 replicates were used for testing as those depicted in Figure 4 (replicates 2, 4, 5, and 7).

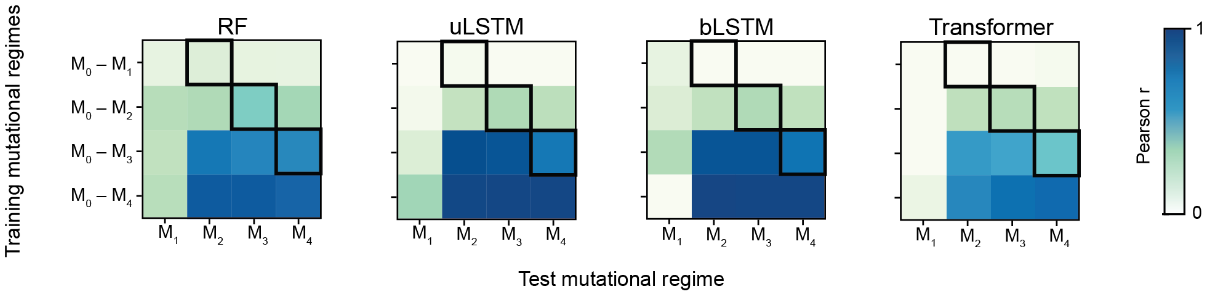


**SI Figure 6. Interpolation and extrapolation performance on GB1.** Heatmaps showing RF, uLSTM, bLSTM and transformer model interpolation and extrapolation performance (Pearson *r*). Cells in a black border represent extrapolation into one mutational regime, to the left is interpolation and to the right extrapolation to higher mutational regimes. The y-axis of the heatmaps corresponds to training mutational regimes, and the x-axis to testing mutational regimes.

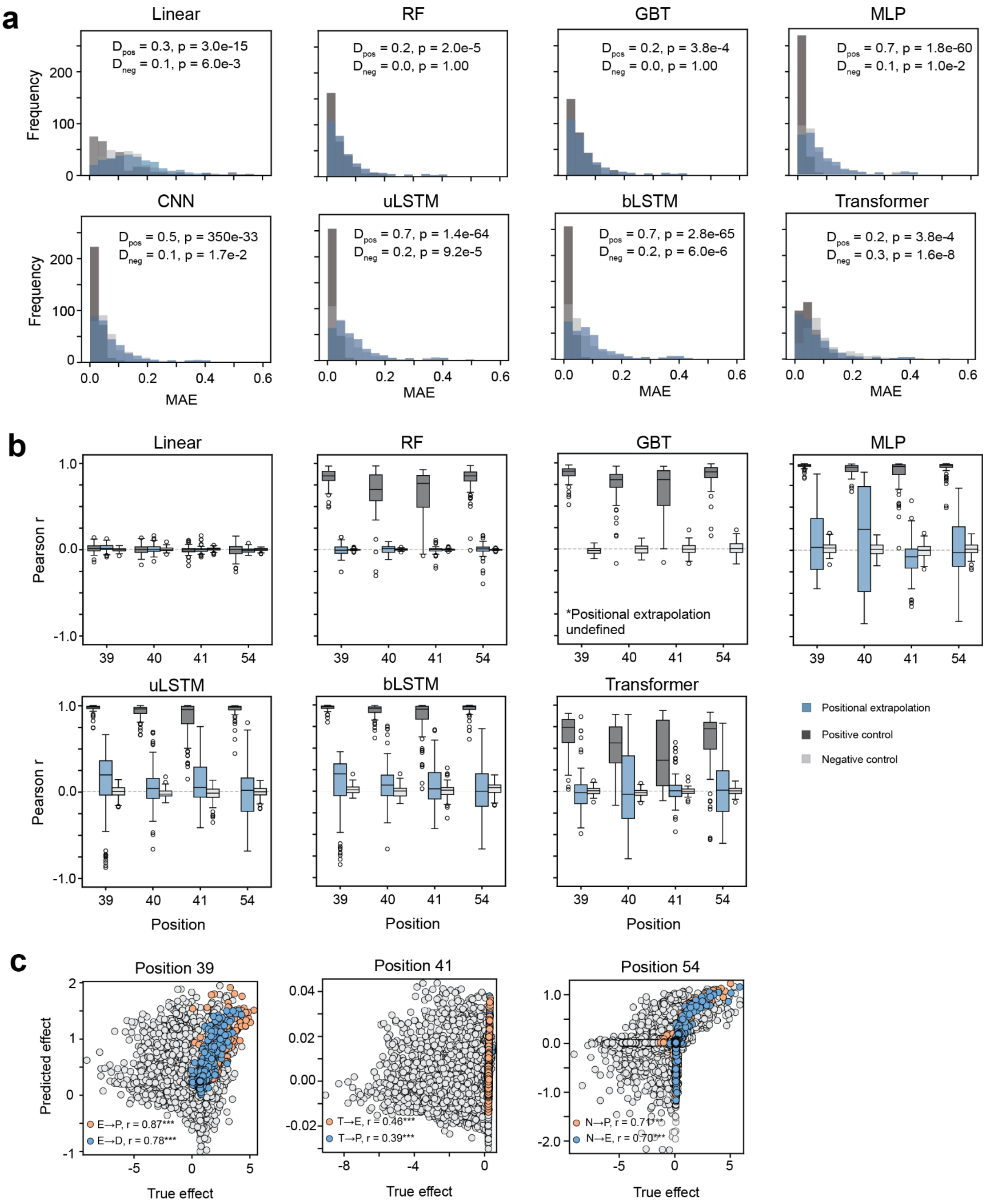


**SI Figure 7. Positional extrapolation on the GB1 dataset.** (**a**) Histogram of MAEs for all models when predicting mutational effects in the positional extrapolation testing regime on the GB1 dataset. D is Kolmogorov's D statistic when comparing the MAEs associated with positional extrapolation against the negative ($D_{neg}$) and positive ($D_{pos}$) control. D is 1 for sample distributions drawn from separate underlying distributions and 0 for those drawn from the same distribution. P-values < 0.05 are considered to be statistically significant. The positive control represents data with the same splitting regime with 80% of sequences containing amino acids other than the fixed amino acid reintroduced to the training dataset and the remaining 20% of sequences used as a test dataset. The negative control represents neural network models with randomly initialized weights or decision tree models trained with shuffled training targets. (**b**) The correlation (Pearson *r*) between predicted mutational effects from linear, RF, GBT,

MLP, uLSTM, bLSTM and transformer models and true mutational effects per position. (**c**) The predicted mutation effects from the CNN model against the true mutational effects for one replicate (replicate 0, seed sequence DETN) at position 39, 41, and 54. Positional extrapolation is separated for amino acid substitutions with the top two Pearson correlation values when comparing predicted and true mutational effects across all observed genetic contexts. Significance is annotated as: *** $p < 0.001$, ** $p < 0.01$, * $p < 0.05$.

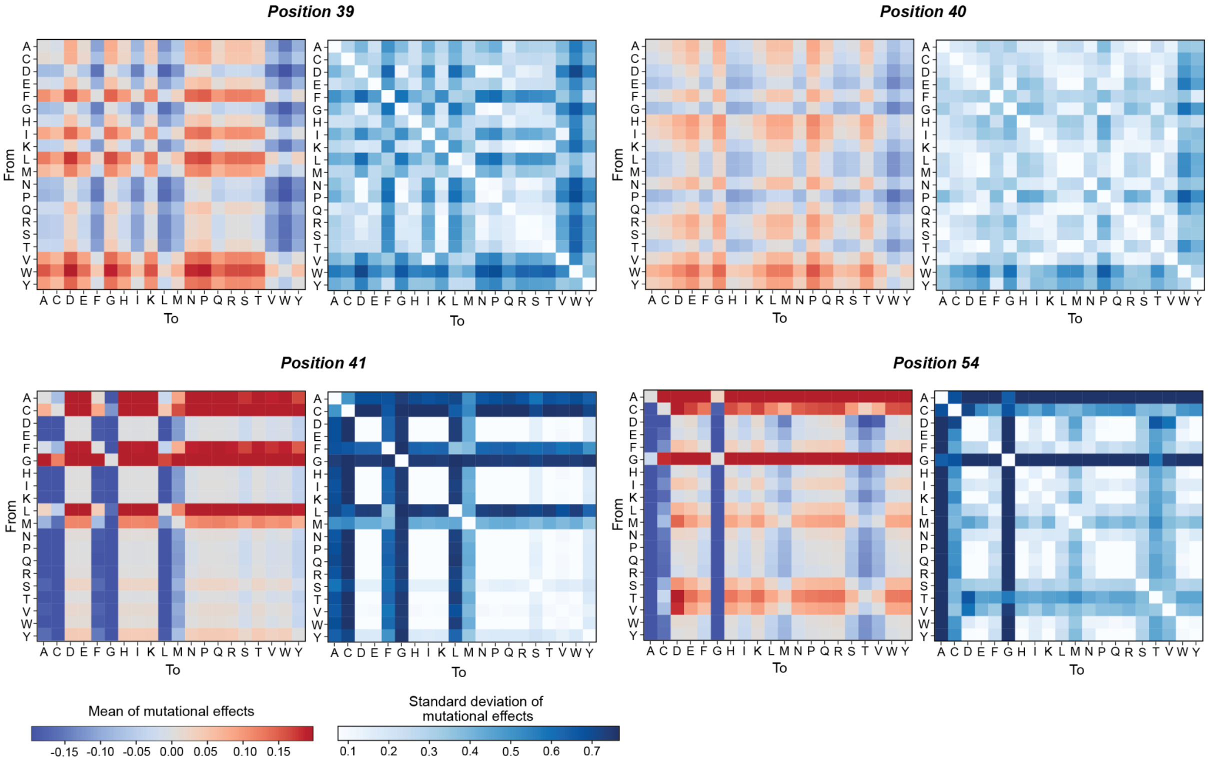


**SI Figure 8. True mutational effects on the GB1 dataset.** The mean (left) and standard deviation (left) of the true mutational effects for each seed amino acid (from) and mutant amino acid (to) at positions 39, 40, 41 and 54.

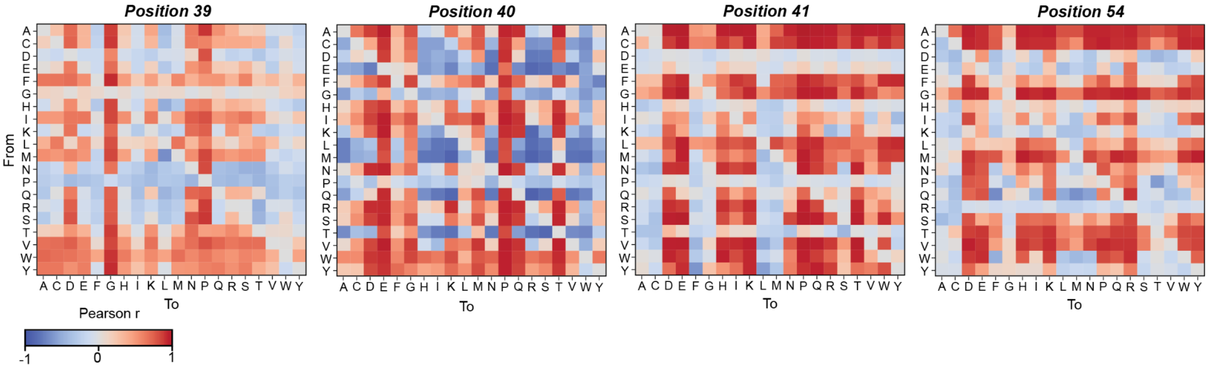


**SI Figure 9. Predicted mutational effects by CNN models on the GB1 dataset.** The predicted mutation effects from the CNN model against the true mutational effects at each position (39, 40, 41, and 54) of the GB1 datasets.

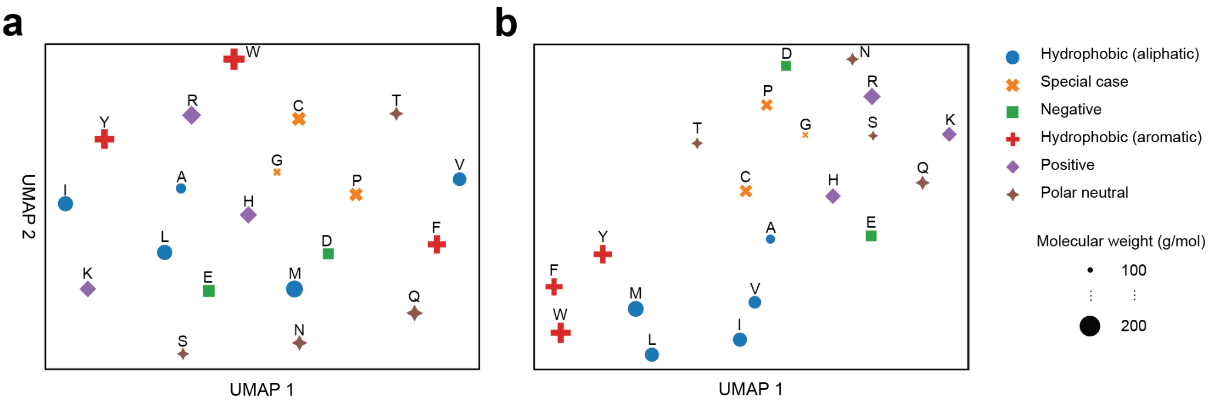


**SI Figure 10. Two-dimensional visualization of amino acid representations.** (a) OHE representations and (b) the final hidden dimension from a trained CNN model were reduced to two-dimensions with UMAP.
